# Genetic Diversity of Bundibugyo Ebolavirus from Uganda and the Democratic Republic of Congo

**DOI:** 10.1101/2021.10.18.464898

**Authors:** Isaac Emmanuel Omara, Sylvia Kiwuwa-Muyingo, Stephen Balinandi, Luke Nyakarahuka, Jocelyn Kiconco, John Timothy Kayiwa, Gerald Mboowa, Daudi Jjingo, Julius J. Lutwama

## Abstract

**Background:** The Ebolavirus is one of the deadliest viral pathogens which was first discovered in the year 1976 during two consecutive outbreaks in the Democratic Republic of Congo and Sudan. Six known strains have been documented. The *Bundibugyo Ebolavirus* in particular first emerged in the year 2007 in Uganda. This outbreak was constituted with 116 human cases and 39 laboratory confirmed deaths. After 5 years, it re-emerged and caused an epidemic for the first time in the Democratic Republic of Congo in the year 2012 as reported by the WHO. Here, 36 human cases with 13 laboratory confirmed deaths were registered. Despite several research studies conducted in the past, there is still scarcity of knowledge available on the genetic diversity of *Bundibugyo Ebolavirus*. We undertook a research project to provide insights into the unique variants of *Bundibugyo Ebolavirus* that circulated in the two epidemics that occurred in Uganda and the Democratic Republic of Congo

**Materials and Methods:** The Bioinformatics approaches used were; Quality Control, Reference Mapping, Variant Calling, Annotation, Multiple Sequence Alignment and Phylogenetic analysis to identify genomic variants as well determine the genetic relatedness between the two epidemics. Overall, we used 41 viral sequences that were retrieved from the publicly available sequence database, which is the National Center for Biotechnology and Information Gen-bank database.

**Results:** Our analysis identified 14,362 unique genomic variants from the two epidemics. The Uganda isolates had 5,740 unique variants, 75 of which had high impacts on the genomes. These were 51 frameshift, 15 stop gained, 5 stop lost, 2 missense, 1 synonymous and 1 stop lost and splice region. Their effects mainly occurred within the L-gene region at reference positions 17705, 11952, 11930 and 11027. For the DRC genomes, 8,622 variant sites were identified. The variants had a modifier effect on the genome occurring at reference positions, 213, 266 and 439. Examples are C213T, A266G and C439T. Phylogenetic reconstruction identified two separate and unique clusters from the two epidemics.

**Conclusion:** Our analysis provided further insights into the genetic diversity of *Bundibugyo Ebolavirus* from the two epidemics. The *Bundibugyo Ebolavirus* strain was genetically diverse with multiple variants. Phylogenetic reconstruction identified two unique variants. This signified an independent spillover event from a natural reservoir, rather a continuation from the ancestral outbreak that initiated the resurgence in DRC in the year 2012. Therefore, the two epidemics were not genetically related.

## Introduction

The Ebolavirus is one of the deadliest viral pathogens which was first discovered in the year 1976 during two consecutive outbreaks in the Democratic Republic of Congo (DRC) and Sudan(1). Since then, over 30 different outbreaks have been reported in Sub-Saharan Africa with an estimated 14,000 deaths and case fatality rates of up to 90% (2)(1). These viruses belong to the family Filoviridae and Genus Ebolavirus (2). There are six known strains in the genus Ebolavirus, all of which have a negative sense-single stranded RNA genome of approximately 18 -19 kilo base pairs (3). They include; *Zaire Ebolavirus*, *Sudan Ebolavirus*, *Bundibugyo Ebolavirus*, *Reston Ebolavirus*, *Tai Forest Ebolavirus* (4) and *Bombali Ebolavirus (5)*. The first three strains have been documented to cause severe illness and death in both humans and non-human primates with case fatality rates ranging from 40%-90% (6) (7). The *Reston* and *Tai Forest Ebolavirus* have not yet been discovered to cause human mortalities(1). Since its first discovery in the year 1976, there has been recurrent outbreaks of Ebolaviruses in Sub Saharan African countries(8)(9)(1)(10). With new cases reported almost every after five years in East and Central Africa for example, Uganda has reported seven different Ebolavirus outbreaks since the year 2000 and the DRC has recorded its 12^th^ outbreak this year in February 2021(1)(11)(12). In particular, the B*undibugyo Ebolavirus* has a genome size of 18,940 base pairs and its RNA genome encodes seven structural proteins namely; Nucleoprotein (NP), two virion proteins (VP35 and VP40), a surface Glycoprotein (GP) and additional two virion proteins (VP30 and VP24). The genome also consists of an RNA- dependent, RNA polymerase (L) and a non-structural soluble protein (sGP proteins). The L gene codes for the RNA Polymerase, which is the most conserved region where as the VP40 virion protein is the most polymorphic gene in the Ebolavirus. The Bundibugyo Ebolavirus made its first appearance on the 1^st^ August 2007, when there were reported cases of a viral hemorrhagic fever in Bundibugyo and Kikyo townships, a district in the western part of Uganda (11). This outbreak resulted into 116 human cases and 39 laboratory confirmed deaths (13). The index case was suspected to be a 26-year-old woman from Kabango village in Bundibugyo district. She presented with general weakness, fever and diarrhea after which she was hospitalized (13). Together with other suspect cases, blood samples were collected, sent to the Uganda Virus Research Institute (UVRI) and the US Centers for Disease Control and Prevention. Several laboratory investigations were performed and they confirmed on the 29^th^ November 2007, a very unique and therefore novel strain of Ebolavirus, that was named Bundibugyo (13) (14). The Epidemiological data collected from this investigation found hunting spears near her home but hunting as a practice was denied.

In order to strengthen the response preparedness of the Viral Hemorrhagic Fever (VHF) in Uganda, the UVRI re-initiated the VHF National Surveillance programme in the year 2010 [34]. This was through an agreement between the Uganda Ministry of Health, Uganda Virus Research Institute and the US Centers for Disease Control and Prevention [32]. To date, UVRI serves as the national and regional reference laboratory for detection and response to VHF outbreaks which are of public health relevance in the region [35]. Currently, there is improved diagnostics to provide real time reporting of VHF cases detected. Laboratory diagnostic assays that have been implemented include; IgM and IgG ELISA for antigen-detection, RT-PCR as well as sequencing [36].

Since then, several research studies have been conducted for example; the work done by J.S. Towner *et al,* 2008 highlighted the high level of genetic diversity at amino acid level in the encoded virus proteins computing to over 27% and 35% for *Bundibugyo* and *Zaire Ebolavirus* respectively (11). Secondly, other research studies done elsewhere have also reported that variations in the Ebolavirus genome might have effects on the efficacy of virus detection at a sequence based level and design of candidate therapeutics (15). Thirdly, several years back, a research study that was conducted following two simultaneous occurrences of Ebolavirus in the DRC and South Sudan in 1976 (16), found a correlation between Ebolavirus disease and animal disease outbreaks (17). This is because Ebolavirus is transmitted by direct contact with the blood or any other secretions from animals or persons (18) (19). In addition, more recent studies that involve the Polymerase Chain Reaction (PCR) and antibody tests have identified cave-dwelling fruit bats as the possible natural reservoirs to most Ebolavirus strains (20) (21). Spillover events therefore occur when the animal and human interface is bridged through human activities such as; hunting wildlife for bush meat (22). This then sparks of epidemics which is most often followed by sustained human to human transmissions (23) (24). Despite all these research studies, five years after the ancestral outbreak was declared over in Uganda, the *Bundibugyo Ebolavirus* re-emerged and caused an epidemic for the first time in the DRC (25). This was reported by the WHO on the 17^th^ August 2012 in Isiro Province (25) (26). The putative index case for this epidemic remains unidentified [14]. However, the earlier laboratory investigations using the RT-PCR assays confirmed, a clinic nurse in Isiro Province whose symptoms began on the 28^th^ June 2012 (25). She reported with multiple potential exposures like human contact with other sick people, exposure to bats and as well she attended a funeral service (25). This outbreak resulted into 36 human cases with 13 laboratory confirmed deaths (26). Despite all these research studies, there is still limited scientific information available to explain the genetic diversity of this strain.

Further to this, in light of new evidence from the February 2021 outbreaks of Ebolavirus in the Republic of Guinea and the DRC, a new and unique paradigm or pattern for how these outbreaks spark off has been identified (27) (17). This new research findings suggests that the putative index cases leading to the resurgences of the February 2021 outbreaks are linked to contacts with survivors from past Ebolavirus outbreaks (28). Surprisingly, the previous outbreak of Ebolavirus in Guinea occurred 5-7 years ago at the time of the West African outbreak (29) (30). Whereas the resurgence in the DRC occurred a year after the 2020 outbreak was declared over (17) This cases have already raised important new research questions such as; “How do we need to change our response to escape from the cycle of outbreak-response-re-introduction-outbreak”, “can new therapeutics be used to clear viruses from survivors” and the immediate question is, what these new findings mean for Ebolavirus survivors who are already faced with a lot of challenges (31). This therefore has created a need for reconsideration into local and scientific accounts of past Ebolavirus outbreaks (27) for example; the two epidemics of *Bundibugyo Ebolavirus* in Uganda in the year 2007 (11) and the DRC in the year 2012 (25). We therefore undertook a research study to get a better understanding of these two epidemics. Our main aim was to determine the genetic diversity of *Bundibugyo Ebolavirus* from Uganda and the Democratic Republic of Congo. The specific goals were to; i) To identify the unique variants in isolates of *Bundibugyo Ebolavirus* from the epidemics that occurred in Uganda and the Democratic Republic of Congo, ii) To determine the genetic relatedness between the *Bundibugyo Ebolavirus* outbreaks in Uganda (2007) and the Democratic Republic of Congo (2012). Ultimately, we aimed to determine whether the resurgence of *Bundibugyo Ebolavirus* in DRC was an independent spillover event from nature or a continuation from the ancestral outbreak, possibly through contacts with past survivors.

**Fig 1:**
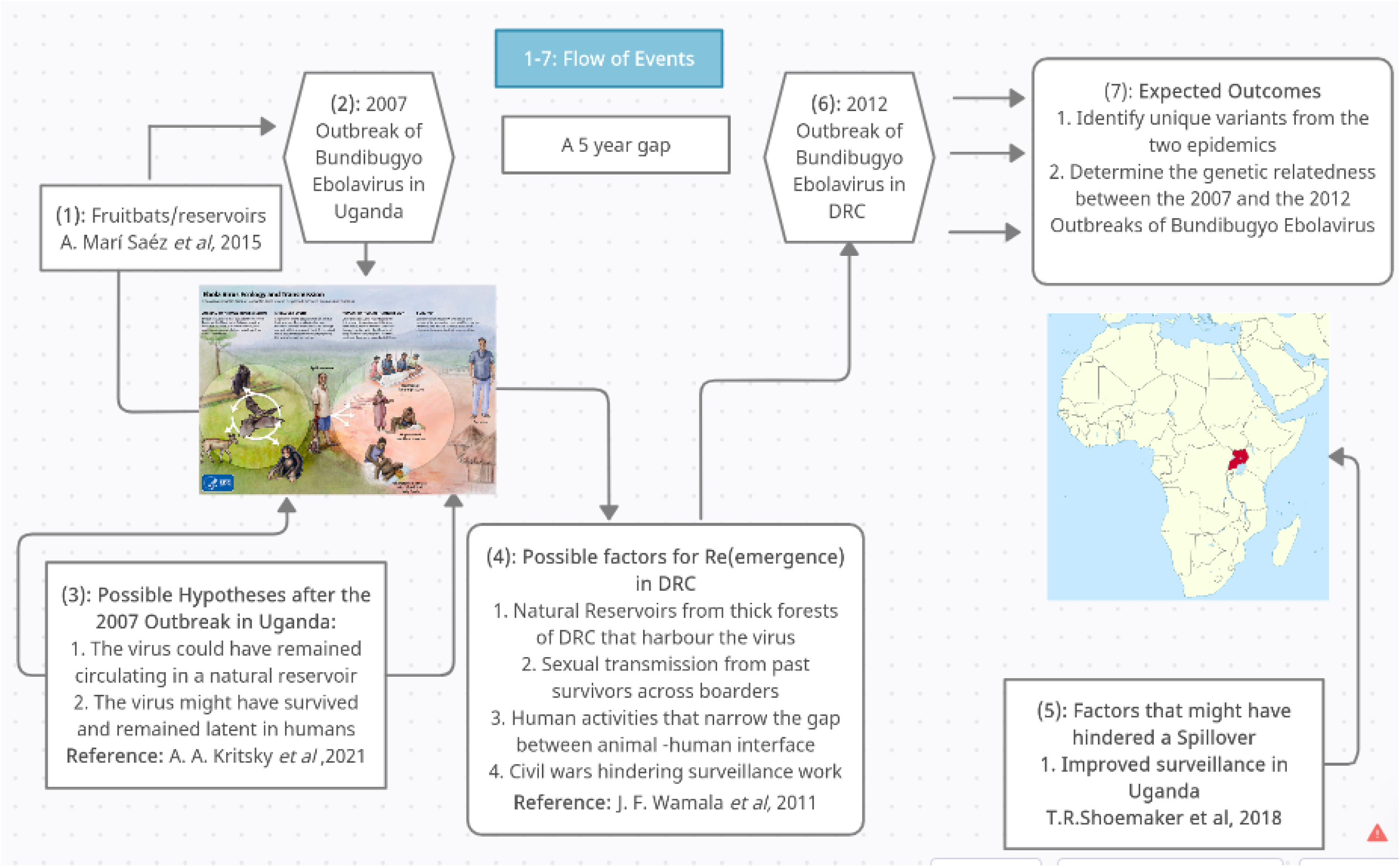
The working hypotheses for the resurgence in DRC in the year 2012.

**Fig 2:**
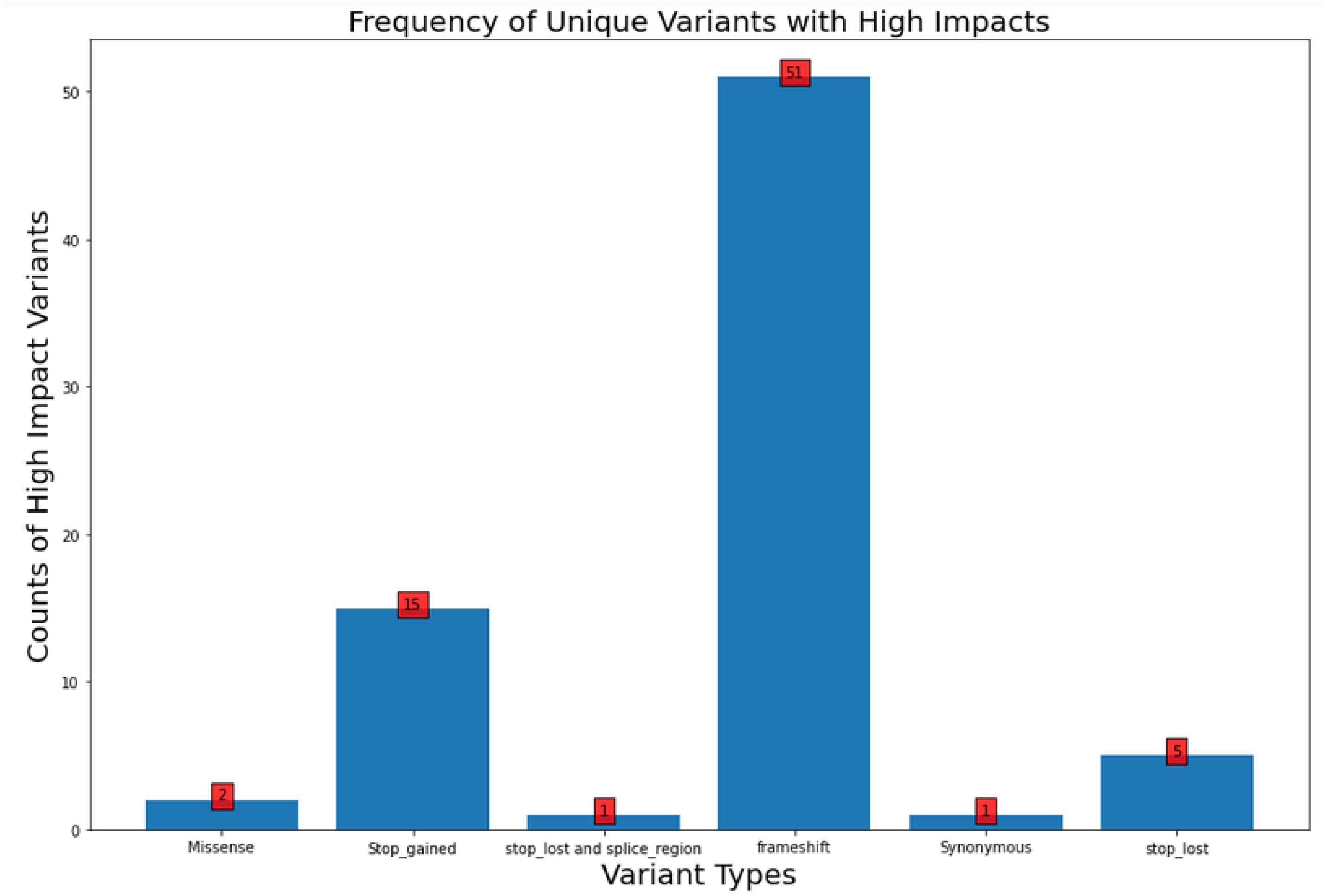
A Bar-plot showing the frequency of unique variants with high impact on the genomes of Bundibugyo Ebolavirus isolated from the 2007 outbreak in Uganda.

## Materials and Methods

### Study Design

This was a retrospective descriptive study. The source organism in our analysis was the *Bundibugyo Ebolavirus* strain, which has a genome size of 18,940 base pairs (32). We used publicly available sequence data that was retrieved from the National Center for Biotechnology and Information Gen-bank database (33) The NCBI, has the Sequence Read Archive (SRA) repository and the Nucleotide sequence database (34). The SRA is the largest publicly available repository having raw sequence data from high throughput sequencers (35). The 31 raw sequence data which represented isolates collected from the epidemic in Uganda was retrieved from this repository. Whereas the Nucleotide sequence database has assembled genomes deposited from different experiments (36). The 4 nucleotide sequences that represented isolates from the DRC in our analysis was retrieved from this database.

### Sample Size Determination

The isolates were; 31 fastq sequences, 6 fasta sequences from the Uganda outbreak in the year 2007. The isolates from the DRC outbreak were represented by the 4 fasta sequences that were retrieved from the nucleotide sequence database (36). The table 1 below shows the characteristics of the isolates which the DRC sequences were generated (25)

**Table 1:** Patient and Sample characteristics from the DRC outbreak in the year 2012.

|  |  |  |  |  |  |
| --- | --- | --- | --- | --- | --- |
| Case ID | Gene-bank Number | Demographics | Occupation | Sampling Location | Clinical Status |
| 112 | KC545393 | 44/F | homemaker | Isiro Province | deceased |
| 120 | KC545394 | 77/M | unknown | Vungba | deceased |
| 122 | KC545395 | Unknown | unknown | Unknown | survived |
| 37 | KC545396 | 18/F | student | Isiro Province | deceased |
| F, female; M, Male |  |  |  |  |  |

### Bioinformatics Analysis

The fastq-dump tool (37) was used to download all the sequence data from the NCBI database. This included the sequences with their metadata which were all stored in a High-Performance Computing (HPC) server at the African Center of Excellence in Bioinformatics and Data Intensive Sciences (38). Quality Assessment was not performed on the DRC sequences. This is because, they were assembled genomes. However for the 31 raw sequence data (fastq format) collected from the Uganda outbreak in the year 2007, a quality control check was performed comprehensively in order to ensure they were of good quality before downstream analysis (39). This assessment was performed using tools; Fast-QC (v0.11.9) (40) and Multi-QC (v1.9) (41) respectively. The low quality regions were then trimmed, including adapter sequences setting the phred threshold at 20 (42).

### Variant Analysis

For the Uganda genomes, the quality filtered raw sequence data were referenced mapped against a reference genome using the Burrow’s Wheeler Aligner tool (0.7.17-r1188) (43) The reference genome used was an isolate from the 2007 outbreak in Uganda (Gene-bank accession number FJ217161). This isolate was used because it has a complete genome size of 18,940 base pairs and it is also an original isolate from the ancestral outbreak. Variants were then called using freebayes tool (v1.3.1-dirty) (44)(45). This tool has advantages over other tools because, it is haplotype based, also a Bayesian genetic variant detector and outputs a variant call format (VCF) file, which consists of small polymorphisms specifically SNPs (single-nucleotide polymorphisms), Indels (Insertions and Deletions), MNPs (multi-nucleotide polymorphisms) (45). We then used SnpEff tool to perform variant annotation (46). This tool predicts the functional effect of the variants on proteins or amino acid changes (47).

To annotate variants, a database from the reference genome has to be built. This was performed using “SnpEff build” tool. To create the SnpEff database, we downloaded sequence data from NCBI for the reference genome of *Bundibugyo Ebolavirus* with accession number, FJ217161. We also downloaded the corresponding General Feature Format (GFF) file, which contains the annotations and the FASTA file, with its entire genome (48) The SnpEff tool was then used to annotate variants. Once the analysis was executed, the annotation data was outputted as an annotated Variant Call Format (VCF) and an HTML report file containing all the summary statistics for the different variants (46). In addition, Python v3.6.3 (49) was then used to construct a bar plot to show the frequency of unique variants with high impacts on the genome.

On the other hand, all the DRC sequences including an isolate of the *Bundibugyo Ebolavirus* from the 2007 outbreak as the reference sequence (Gene-bank accession number FJ217161) were concatenated in to a single multi-fasta file and saved as a FASTA format. This reference sequence was used in order to determine how the variants from the 2012 outbreak were phylogenetically distinct from the 2007 outbreak in Uganda. Multiple Sequence Alignment was performed on the fasta sequences using MAFFT v7.310 tool (50). After this, variants were then called using the alignment FASTA file as input and the SNP extraction tool, SNP-sites v2.3.3 (51). This tool restructures the aligned data as a Variant Call Format (VCF) file. This VCF file provides a clear mapping of SNPs from the aligned sequences. This then allowed easy identification of the SNP location and the genotype for each sample at a given locus (52). In the outputted VCF file, the rows correspond with each unique variant and the columns provides the genotype at that given site (53). A summary of the SNPs relative to the reference sequence was then visualized using the snipit tool (https://github.com/aineniamh/snipit) and SnpEff tool was used to annotate the variants. Using different bash scripts, a report showing the effect of the variants on the different sequences was extracted (54)

### Phylogenetic Analysis to determine the genetic relatedness between the two Epidemics

All the quality filtered raw sequence data from Uganda, were assembled using both SPades v3.13.1 (55) and abyss 1.9.0 assemblers (56) (57) in order to obtain a consensus sequence. These two genome assemblers are best suited for assembly of short paired end reads (58). A draft scaffold was then obtained with the use of SSPACE tool (59). This is a standalone tool and was used for scaffolding the paired end reads. It enabled read orientation into connected sequences by allowing mean values and standard deviations of the insert sizes for each read library (59). GapFiller tool was then used to find and fill gaps generated in the contiguous sequence (60). All the obtained fasta sequences from both Uganda and the DRC were concatenated in to a single FASTA file including the reference sequence. Multiple Sequence Alignment was then performed using MAFFT v7.310 (50) and manually checked in Ali-view v.1.27(61). The 5’ and the 3’ untranslated regions were then trimmed to remove any remaining gaps. The maximum-likelihood phylogenetic tree was then constructed using IQ-TREE (62) and Phyml (63) and the best suited substitution model was determined and run for 1000 replicates. The resulting newick file was uploaded to the interactive tree of life, iTOL v4.0(64), which is an online tool for phylogenetic tree visualization. The tree was rooted at mid-point to split variants from Uganda and the Democratic Republic of Congo.

## RESULTS

### Variant Analysis

In the Uganda genomes, 37 sequences of *Bundibugyo Ebolavirus* were analyzed. We identified 5,740 distinct genome variants and they are recorded in table 2 below. Generally, the variants were distributed according to the different regions (downstream, upstream, exon intergenic, 3 and 5 prime Untranslated Regions (UTR). However, majority of the variants were found downstream and upstream regions of the genome (non-coding regions). The viral sequences showed multiple diversity with most variants occurring at reference positions 17705, 11952, 11930 and 11027 appearing in most of the isolates collected from this outbreak.

**Table 2:**
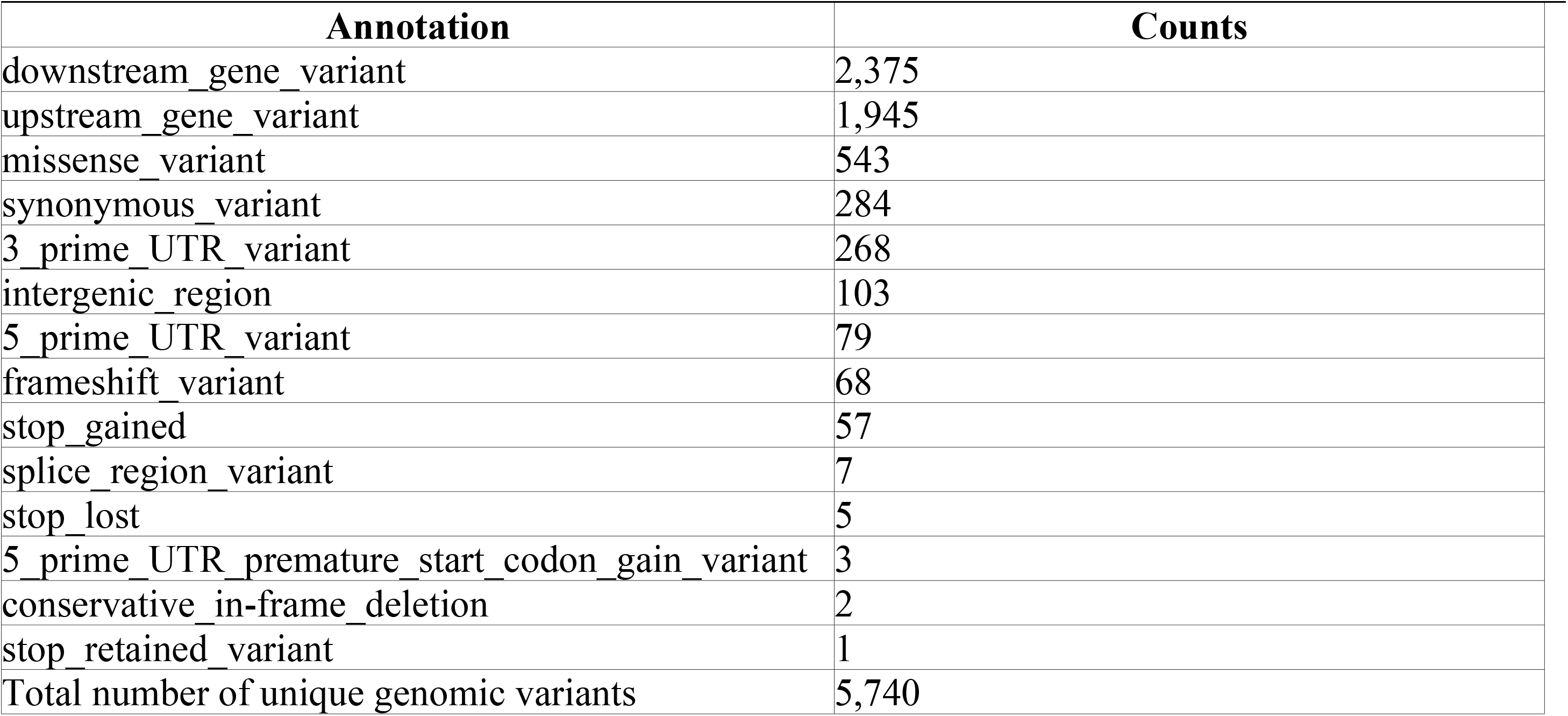
The frequency of each unique variant type in isolates of the *Bundibugyo Ebolavirus* collected from the 2007 outbreak in Uganda.

| Number of Effects by Type |  |
| --- | --- |
| Annotation | Counts |
| downstream gene variant | 2,375 |
| upstream gene variant | 1,945 |
| missense variant | 543 |
| synonymous variant | 284 |
| 3_prime_UTR_variant | 268 |
| intergenic_region | 103 |
| 5_prime_UTR_variant | 79 |
| frameshift variant | 68 |
| stop_gained | 57 |
| splice_region_variant | 7 |
| stop_lost | 5 |
| 5_prime_UTR_premature_start_codon_gain_variant | 3 |
| conservative_in-frame_deletion | 2 |
| stop_retained_variant | 1 |
| Total number of unique genomic variants | 5,740 |

In addition, we then identified 75 unique variants of high impacts on the genome. They were; 51 frameshift, 15 stop gained, 5 stop lost, 2 missense, 1 synonymous and 1 stop lost and splice region. Among these variants, the most common impacts were majorly frame-shifts and stop gained. They include; T**A**T17705TT, GAAAAAATTTTG11952GAAAAAA**A**TTTTG, G11930T, CAAAAAACCCG11027CAAAAAA**A**CCCG. Their effects occurred mostly on the L-gene region of the *Bundibugyo Ebolavirus*. Refer to S2 in Appendix for the supporting information showing a table indicating the frequency of unique variants which had high impacts on the genome.

On the other hand, we identified 8,622 nucleotide variant sites from the isolates of *Bundibugyo Ebolavirus* collected from the 2012 outbreak. The variants identified here all had a modifier effect on the genome. This effect was predetermined by the variant type in each of the isolates. Some of them include; C213T, A266G and C439T. Fig 3 below shows the different nucleotide variant sites in the DRC sequences relative to the reference sequence with Gene-bank accession number of FJ217161. This was a complete genome and isolated from the 2007 outbreak. The purpose of using this as a reference sequence was to find out how the sequences from the 2012 outbreak in the DRC were genetically distinct and unique from the ancestral outbreak of 2007 which occurred in Uganda.

**Fig 3:**
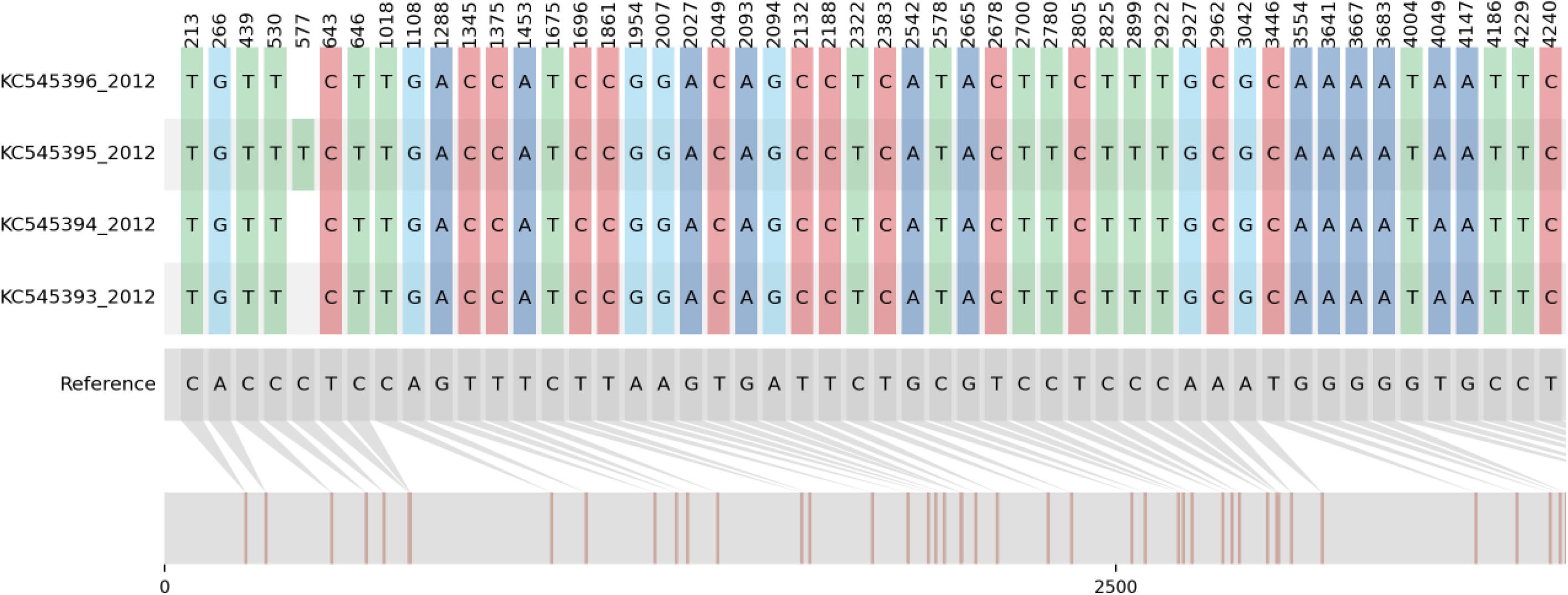
Nucleotide alignment showing variant sites in the four sequences relative to the reference genome. The 4 sequences represent isolates collected from the DRC outbreak in 2012 (NCBI accession numbers: KC545393, KC545394, KC545395, KC545396). The reference sequence is an isolate from the ancestral outbreak (Gene-bank accession number: FJ217161)

### Phylogenetic Analysis to determine the genetic relatedness between the two Epidemics

When the tree in Fig 4 below was rooted at mid-point, two separate and unique clusters were identified from these two epidemics. Phylogenetic reconstruction demonstrates that the 4 sequences isolated from the outbreak in DRC cluster uniquely and distant from those of the 2007 outbreak in Uganda. This signify a separate variant and basing on our analysis, we identified approximately 8,622 mutations from the 4 DRC sequences, which is almost double the number of mutations identified from the ancestral outbreak in Uganda (5,740)

**Fig 4:**
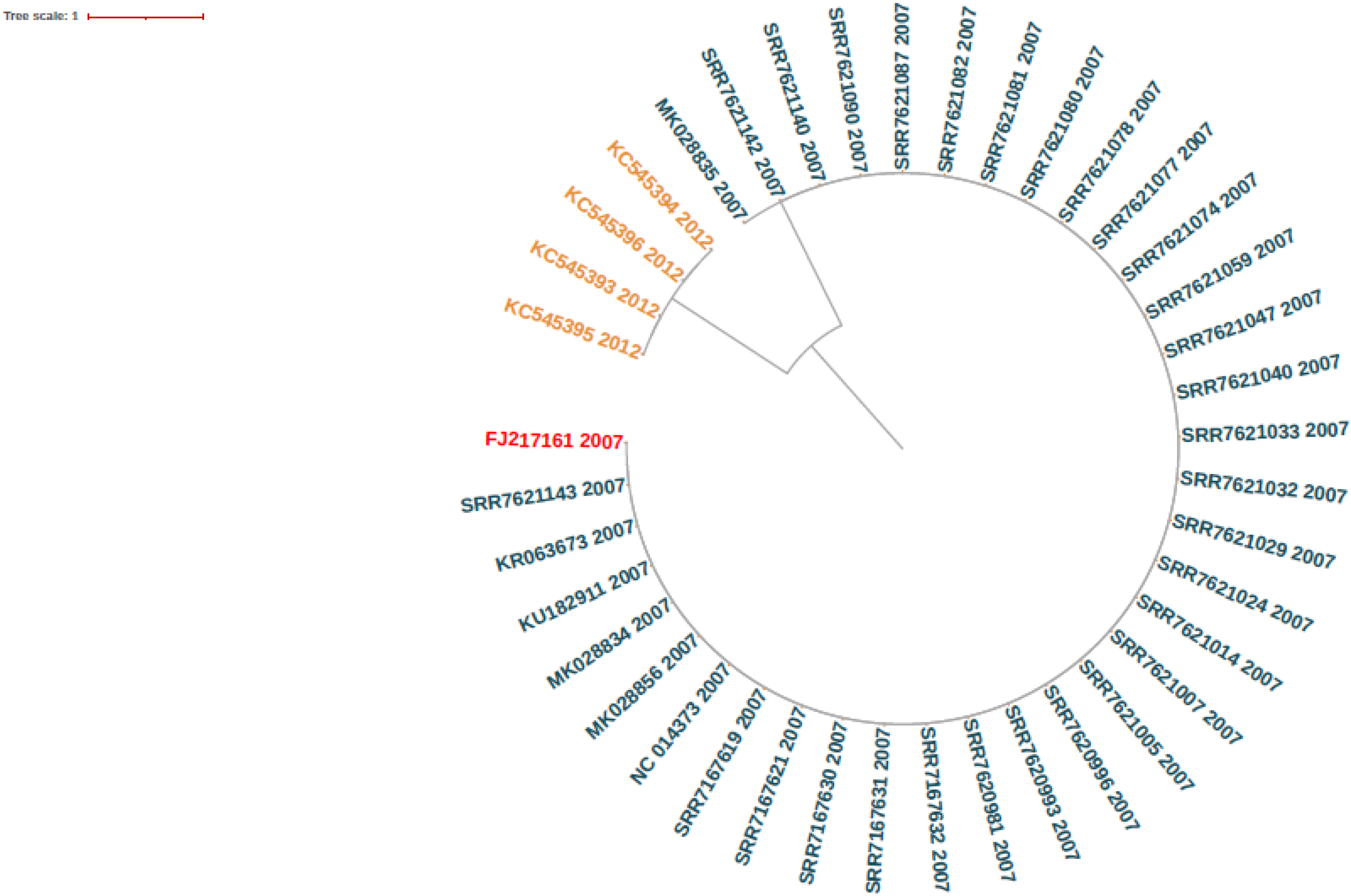
Maximum likelihood phylogenetic tree showing the two unique clusters identified from the two epidemics that occurred in Uganda (2007) and the Democratic Republic of Congo (2012). The reference sequence used was an isolate from the ancestral outbreak having a Gene-bank accession number of FJ217161

## Discussion

We generally aimed to determine the genetic diversity of *Bundibugyo Ebolavirus* from the epidemics that occurred in Uganda in the year 2007 and the DRC in the year 2012. Our specific goal was to identify the unique variants in isolates collected from these two epidemics. This was to ultimately enable us determine the genetic relatedness between two epidemics.

Our analysis identified in total, 14,362 unique genomic variants from the two epidemics. The high impacts variants in the Uganda genomes were mainly frame shifts, stop gained and missense mutations. A frameshift is a genetic variant that changes the way codons are read during the process of creating an amino acid sequence (65). This variant is due to an insertion or deletion of a nucleotide (66). This is of significance because cells read a gene in groups of three bases. The three bases here correspond to one of 20 different amino acids that is used to build a protein (67). Therefore, if a mutation disrupts this reading frame, then the sequence of DNA that follows the mutation will be read incorrectly (68). With a stop gained variant, the mutation leads to changes of at least one base of a codon, hence a premature stop codon (69). This results in a premature stop of translation of messenger RNA in to a protein hence a non-functional or unstable protein (52). In addition, a missense mutation is a genetic change where a single pair of substitution alters the genetic code leading to production of a new amino acid (70). Most of these variant effects occurred within the L-gene region of the *Bundibugyo Ebolavirus*. Since the L-gene is the most conserved region and is a target for primer designs (71), the findings from our analysis is in line with a previous study conducted after the 2007 outbreak (11). Where sequence analysis of the PCR fragment from the virus L-gene revealed the initial failure of real-time RT-PCR assays, since the viral sequences were divergent from the four already known strains of Ebolavirus (11). Therefore, these alterations of the genetic code or disruption of one reading frame could have resulted in to the formation of a new strain of Ebolavirus and hence our findings supports this past study. The high frequency of frameshift variants is suggestive of a new strain (52), the *Bundibugyo Ebolavirus,* which was a novel strain that first emerged in the year 2007 in Uganda (11).

On the other hand, 8,622 nucleotide variant sites were identified from the DRC genomes. The variants had a modifier effect on the genome. This effect was predetermined by the type of variant identified in these isolates (52). Modifier variants are genes that alter the phenotypic outcomes and results in to altered effects or impacts (72). The phenotypic outcomes here could include; dominance, expression and penetrance (73). Naturally, viruses accumulate mutations over time which may arise from adaptations in response to environmental changes or immune responses of the host reservoirs (74). Sometimes viruses transmit and persists after fixing beneficial mutations that would allow it to better exploit it’s host or other new hosts (75). This scenario could explain the re-emergence of the *Bundibugyo Ebolavirus*. That is, after the ancestral outbreak was declared over in the year 2007, this virus might have undergone an evolutionary change over a period of five years in a certain natural reservoir, generating variations in it’s genome. This resulted into a separate and unique variant that was responsible for the 2012 outbreak in the DRC (76). The ultimate high frequency of modifier effects on the genome is an indicator and possibly explains, the divergence or formation of a new variant that was unique from the ancestral type (25)(26).

Phylogenetic trees help in our understanding of the evolutionary relationships between groups (77). In our context, we used it to determine the genetic relatedness between the epidemics that occurred in Uganda in the year 2007 and the DRC in the year 2012. Phylogenetic reconstruction in Fig 4 demonstrates that the 4 sequences from the 2012 outbreak in DRC cluster together and are similar but distantly related from those of the ancestral outbreak (78). This signify a new variant and basing on our analysis, we identified approximately 8,622 mutations from the 4 DRC sequences, which is almost double the number of mutations identified from the ancestral outbreak in Uganda (5,740) (79). This is indicative of viral evolution over the period of five years (80). In other words, the frequency of SNPs or mutations occurrence in a genome under the conditions of a survivor organism is reduced by a big magnitude compared to that from a host reservoir (81). This is because the virus under goes a period of latency in a human survivor (82). Therefore, these two separate variants indicate that the 2012 outbreak in DRC was a new introduction or an independent spillover event from a certain animal reservoir, rather a human transmission from a contact with a past survivor.

Our study however had limitations such as; limited sampling which led to less sequence data generated from the 2012 outbreak. Some patient demographics were unknown, this hindered our understanding in to the Epidemiology and Molecular findings. This led to uncertainty in drawing conclusions on the genetic diversity of *Bundibugyo Ebolavirus* from the 2012 outbreak in the DRC. For example; in variant analysis and phylogenetic estimations. The availability of more or complete genomes from the DRC outbreak in 2012 would improve the study of transmission dynamics between these two epidemics as well as identification of multiple key SNP’s that can promote the study of *Bundibugyo Ebolavirus* pathogenesis.

## In conclusion

our analysis provided further insights into the genetic diversity of Bundibugyo Ebolavirus from the two epidemics. Variant characterization can be used in the fight against Bundibugyo Ebolavirus and the development of effective treatments or vaccines. This is because key SNPs have been identified and can be used for further research about the pathogenesis of *Bundibugyo Ebolavirus.* The findings from our study has also provided knowledge on the likely origin or how the 2012 outbreak in the DRC was initiated. Phylogenetic reconstruction identified two unique variants. This signified an independent spillover event from a natural reservoir, rather a continuation from the ancestral outbreak that initiated the resurgence in the DRC in the year 2012. Therefore, the two epidemics are not genetically related.

## Abbreviations

BDBV: Bundibugyo Ebolavirus
RT-PCR: Reverse Transcription Polymerase Chain Reaction
DRC: Democratic Republic of Congo
SNP: Single Nucleotide Polymorphism
IgM: Immunoglobulin M
IgG: Immunoglobulin G
UVRI: Uganda Virus Research Institute

## Acknowledgments

I would like to extend my sincere gratitude to the department of Immunology and Molecular Biology, College of Health Sciences in Makerere University, for the training leading to the award of a Master of Science in Bioinformatics. Special thanks to the department of Arbovirology, Emerging and Re-emerging Infectious Diseases at the Uganda Virus Research Institute. They offered financial support to facilitate my studies. Finally, I would like to extend my appreciation to the MRC/UVRI and LSHTM Uganda Research Unit for an offer of a Manuscript Mentorship Programme that eventually facilitated the submission of this master’s research project work for publication.

## Supporting Information

S1 Appendix: List of the Gene-bank identifiers for the sequences that were used in our analysis

S2 Appendix: The frequency of unique variants which had high impact on the genomes

## Ethical Clearance

This research project was approved by the School of Biomedical Sciences Research and Ethics Committee (SBSREC). This is an institutional review board found within the College of Health Sciences in Makerere University. The protocol number was SBS-2021-64

## Data Availability

There was no funding for this project. We used publicly available sequence data that was retrieved from the National Center for Biotechnology and Information (NCBI). Below are the links. Raw sequence data: https://www.ncbi.nlm.nih.gov/sra/?term=Bundibugyo+Ebolavirus+in+Uganda Assembled genomes: https://www.ncbi.nlm.nih.gov/nuccore/?term=Bundibugyo+Ebolavirus

## Supporting Information

**S1 Appendix:** List of the Gene-bank identifiers for the sequences that were used in our analysis

| Gene-bank ID's for the Uganda outbreak in 2007 |  | Gene-bank ID's for DRC outbreak in 2012 |
| --- | --- | --- |
| SRR7621005 | SRR7621144 | KC545393 |
| SRR7621077 |  |  |
| SRR7621007 | SRR7167631 | KC545394 |
| SRR7621078 |  |  |
| SRR7621014 | SRR7620993 | KC545395 |
| SRR7621080 |  |  |
| SRR7621024 | SRR7620996 | KC545396 |
| SRR7621081 |  |  |
| SRR7621029 | NC_014373 |  |
| SRR7621082 |  |  |
| SRR7621032 | MK028856 |  |
| SRR7621087 |  |  |
| SRR7621040 | MK028834 |  |
| SRR7621090 |  |  |
| SRR7621047 | KU182911 |  |
| SRR7621140 |  |  |
| SRR7621059 | KR063673 |  |
| SRR7621142 |  |  |
| SRR7621074 | MK028835 |  |
| SRR7621143 |  |  |

**S2 Appendix:**
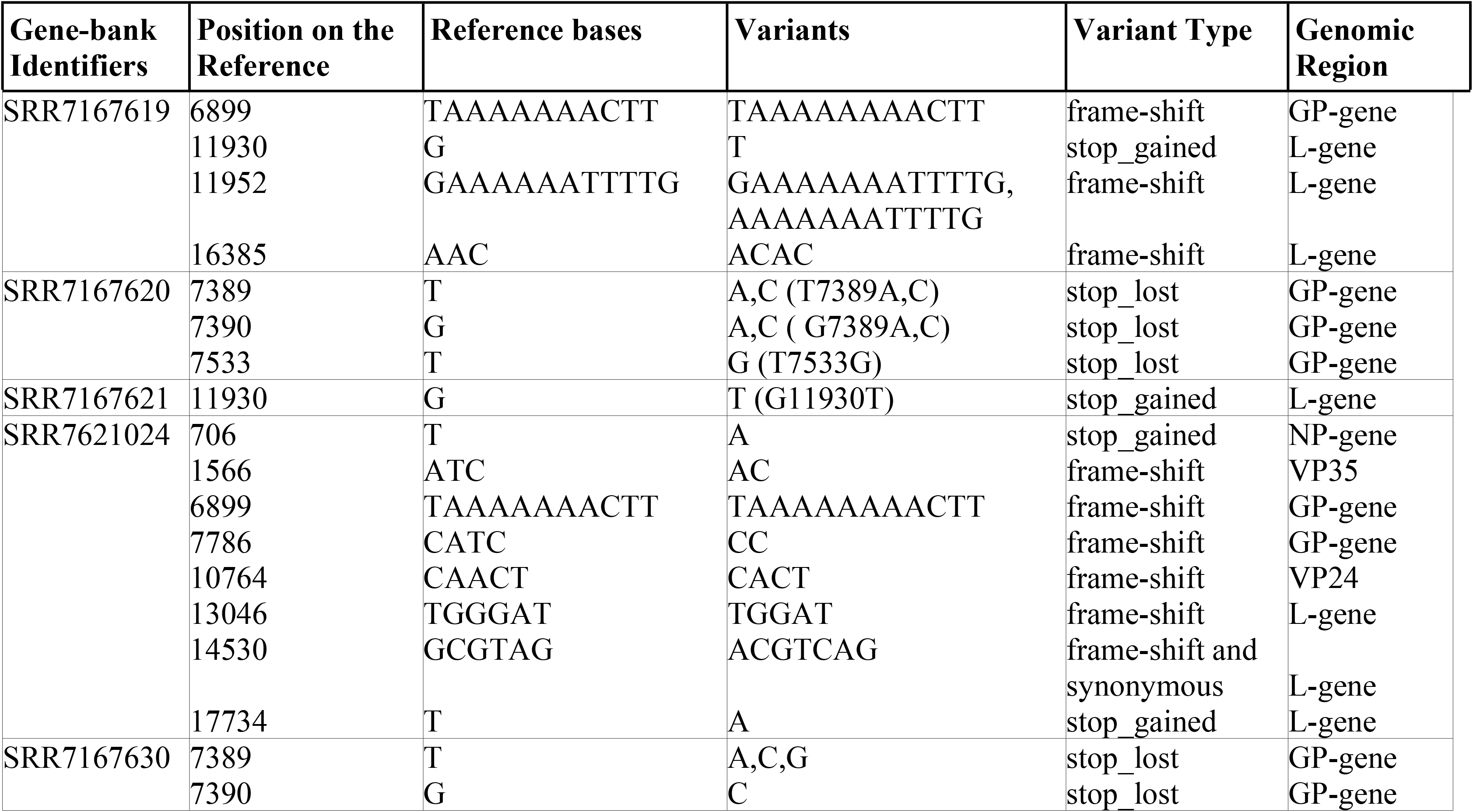
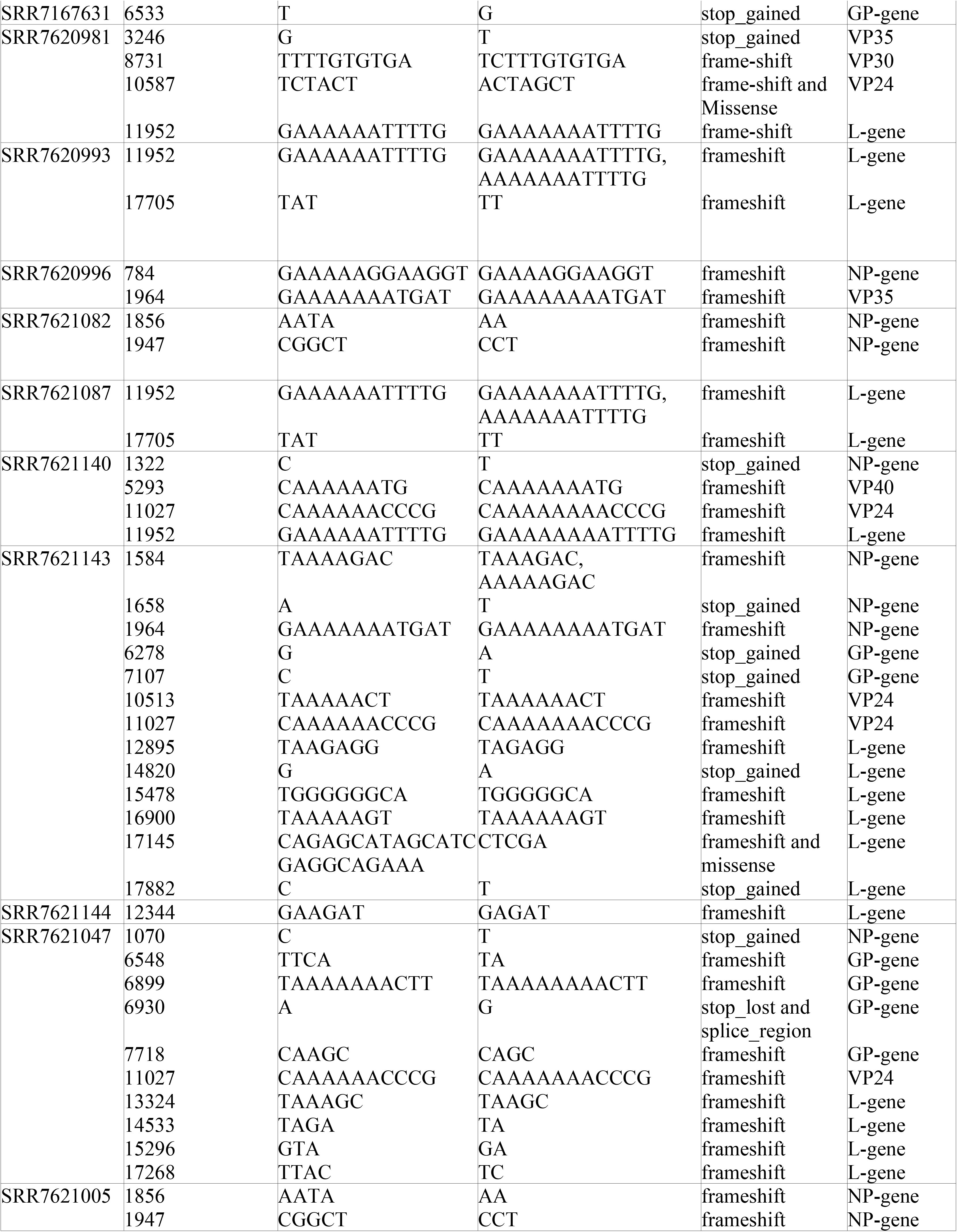
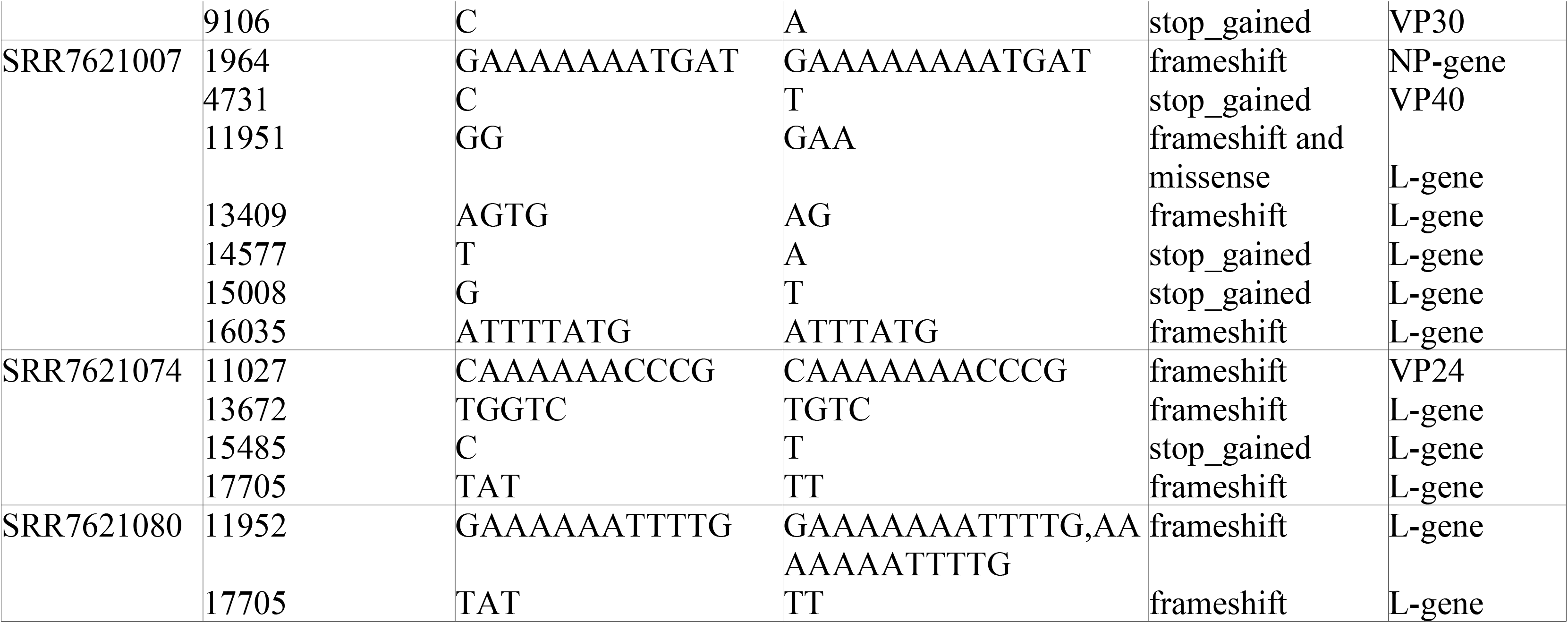
The frequency of unique variants which had high impact on the genomes

## Notes

### Competing Interest Statement

The authors have declared no competing interest.

